# On Classification and Taxonomy of Coronaviruses (Riboviria, Nidovirales, Coronaviridae) with special focus on severe acute respiratory syndrome-related coronavirus 2 (SARS-Cov-2)

**DOI:** 10.1101/2020.10.17.343749

**Authors:** Evgeny V. Mavrodiev, Melinda L. Tursky, Nicholas E. Mavrodiev, Malte C. Ebach, David M. Williams

## Abstract

Coronaviruses are highly pathogenic and therefore important human and veterinary pathogens viruses worldwide (*1*). Members of family Coronaviridae have previously been analysed phylogenetically, resulting in proposals of virus interrelationships (*2–5*). However, available Coronavirus phylogenies remain unrooted, based on limited sampling, and normally depend on a single method (*2–11*). The main subjects of this study are the taxonomy and systematics of coronaviruses and our goal is to build the first natural classification of Coronaviridae using several methods of cladistic analyses (*12*), Maximum Likelihood method, as well as rigorous taxonomic sampling, making the most accurate representation of Coronaviridae’s relationships to date. Nomenclature recommendations to help effectively incorporate principles of binary nomenclature into Coronaviridae taxonomy are provided. We have stressed that no member of *Sarbecovirus* clade is an ancestor of SARS-Cov-2, and humans are the only known host.

**One Sentence Summary:** Multiple comprehensive phylogenetic analyses of all coronavirus species enabled testing of critical proposals on virus interrelationships.

## Introduction

Coronaviridae is a family of the order Nidovirales (realm Riboviria) (*13, 14*). Accordingto the current summaries of International Committee on Taxonomy of Viruses (ICTV) (*13–16*) this family is divided into two subfamilies - Letovirinae and Orthocoronavirinae. These two subfamilies circumscribe five genera: monotypic genus *Alphaletovirus* of the subfamily Letovirinae (with single species *Microhyla letovirus* 1), and four non-monotypic genera of subfamily Orthocoronavirinae, namely:

1. genus *Alphacoronavirus* with 12 subgenera: *Colacovirus* (monotypic), *Decacovirus* (two species), *Duvinacovirus* (monotypic), *Luchacovirus* (monotypic), *Minacovirus* (two species), *Minunacovirus* (two species), *Myotacovirus* (monotypic), *Nyctacovirus* (monotypic), *Pedacovirus* (two species), *Rhinacovirus* (monotypic), *Setracovirus* (two species) and *Tegacovirus* (monotypic) (Table S1);
2. genus *Betacoronavirus* with five subgenera: *Embecoviru*s (four species), *Hibecovirus* (monotypic), *Merbecovirus* (four species), *Nobecovirus* (two species) and *Sarbecovirus* (the number of species is under review) (Table S1);
3. genus *Gammacoronavirus* with two monotypic subgenera: *Cegacovirus* and *Igacovirus* (Table S1);
4. genus *Deltacoronavirus* with three monotypic subgenera *Andecovirus*, *Herdecovirus* and *Moordecovirus* and one non-monotypic subgenus *Buldecovirus* (four species) (Table S1).

The viruses of Coronaviridae, such as the severe acute respiratory syndrome (SARS) and related Middle East respiratory syndrome (MERS), are highly pathogenic (*1*). The recently discovered virus SARS-Cov-2, which is also a member of this family, causes the Coronavirus disease 2019 (COVID-19) that was declared a pandemic by WHO in March 2020. Eight months later the total cases of COVID-19 are approaching 40,100,000 and cumulative deaths have exceeded 1 127 000 (www.who.int, October 2020). Thus, Coronaviridae is of high medical importance worldwide. Such importance has resulted in an urgency to further understand the relationships within the coronavirus family, and the viruses most closely related to SARS-Cov-2. However, phylogenetic analyses of Coronaviridae to date have remained in question due to the use of limited or arbitrary taxonomic sampling, and/or the use of unrooted trees (for example (*2, 3, 6, 8–11*), among others). These issues can result in bias and lack vital hierarchical information needed to understand the critical relationships between the viruses within this family. To address this urgent need, this study has undertaken a comprehensive taxonomic analysis of the family Coronaviridae, using the best taxonomic sampling that is based on all ICTV-approved genomes of all known coronaviruses. Additional viruses were also included to enable testing of the veracity of recently published proposals, included recently identified viruses such as SARS-Cov-2.

Almost all available phylogenetic studies of coronaviruses have been based on the parametric Maximum Likelihood (ML) method. Thus, in order to increase the accuracy of our current analyses and resolve the relationships within Coronaviridae, we have used ML method as well as non-parametric cladistic approaches:

1. standard maximum parsimony (MP),
2. three-taxon statement analysis (3TA), and the
3. Average Consensus (AC) analysis as applied to the array of the maximal relationships.

The last two methods have not previously been used to resolve relationships within the Coronaviridae.

The produced trees (Figure 1, Figures S1 – S5) enabled testing for monophyly of all available non-monotypic genera and infrageneric taxa of Coronaviridae to determine whether they are representative of a natural hierarchy. If so, this validates the estimation of the relationships between all of the available genera/infrageneric taxa or particular virus species of Coronaviridae (incl. SARS-Cov-2).

**Figure 1.**
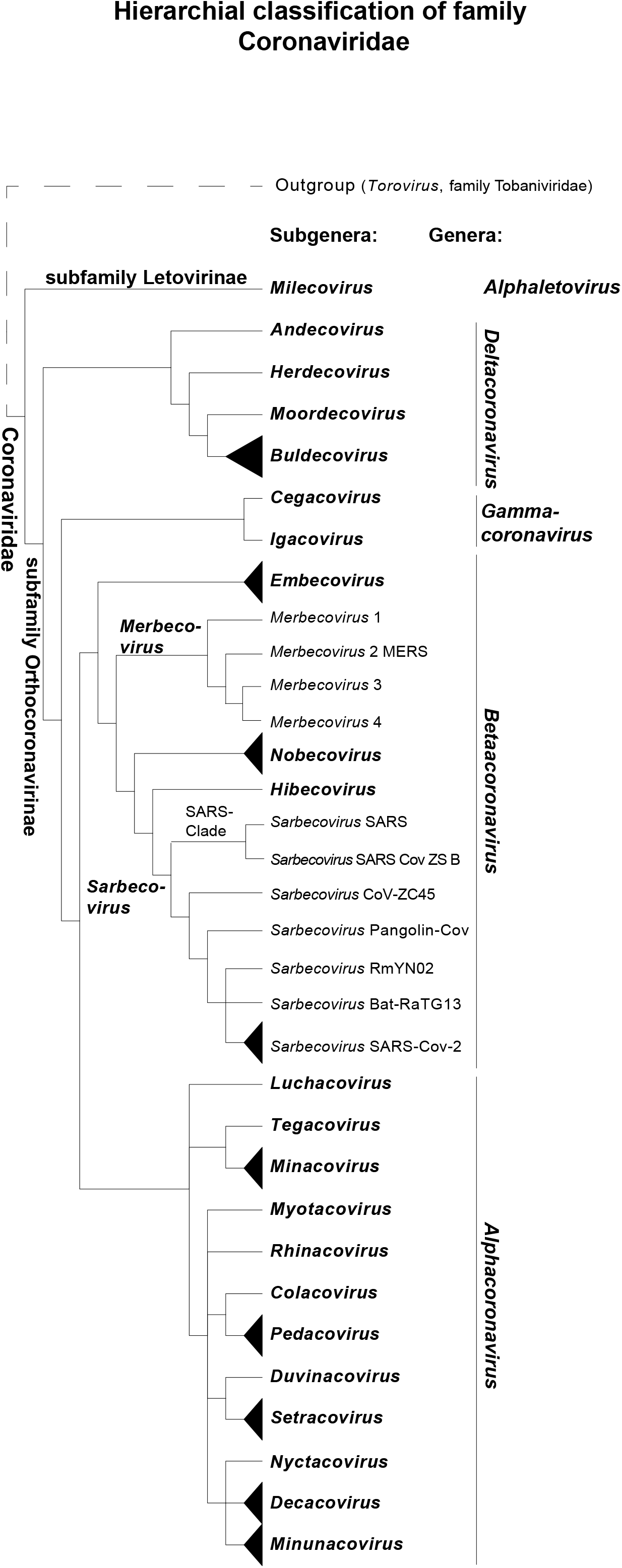
Hierarchical classification of the coronaviruses (Riboviria, Nidovirales, Coronaviridae), established as a simplified Strict Consensus of four trees, produced by three different methods of cladistic analysis as well as by Maximum Likelihood method using G-block version of the genomic alignment of Coronaviridae + *Torovirus*. See Methods (Supplementary Materials), Figures S1 – S5 and Table S1 for more detail including the tree node support values. Figure was drawn with special attention to the taxonomic placements of highly pathogenic Middle East respiratory syndrome-related coronavirus (MERS-Cov) and severe acute respiratory syndrome-related coronaviruses 1 and 2 (SARS-CoV and SARS-Cov-2).

The first important proposal assessed herein is that the newly described monotypic genus *Alphaletovirus* (Table S1), a member of subfamily Letovirinae, is a sister group of family Coronaviridae (*2*).

The second proposal assessed was by Tokarz *et al*. (*4*) who suggested that the general relationship within Coronaviridae subfamily Orthocoronavirinae is a simple hierarchy: ((({*Alphacoronavirus*},{*Betacoronavirus*}){*Gammacoronavirus*}){*Deltacoronavirus*}). Unfortunately Tokarz *et al*. (*4*) based their suggestion on limited taxonomic sampling. We would name this proposition the (((A, B) Γ) Δ) hypothesis of the general relationship within the subfamily.

Thirdly, the subgenus *Hibecovirus* has been placed as sister taxon of the presumably monotypic subgenus *Sarbecovirus* (for instance (*3, 5*)). This proposal is again based on unrooted phylogenies, so it has been tested herein with special attention to the placement of SARS-Cov-2.

In the fourth proposal, the close relationship of SARS-Cov-2 with bat coronaviruses RaTG13 (*21, 25*) and RmYN02 (*24, 25*) as well as with pangolin coronavirus (reviewed in (*17*)) is also tested.

In addition, within the context of our findings, some critical comments are made to the nomenclature of the family as well as to the general patterns of hosting within Coronaviridae, including newly discovered coronavirus SARS-Cov-2.

## Results

All four trees have several major consistencies, as shown in the Strict Consensus (Figure 1 and Figure S5, see also Figures S1 – S4).

### a. General pattern of the relationships within Coronaviridae and the placement of *Microhyla letovirus* 1 (Figure 1, Figure S1 – S5, Table S1)

The results of all analyzes have demonstrated that the hierarchy ((({*Alphacoronavirus*}, {*Betacoronavirus*}) {*GGammacoronavirus*}){.*Deltacoronavirus*}), with *Microhyla letovirus* 1 (subgenus *Milecovirus*, genus *Alphaletovirus*, subfamily Letovirinae) which has been defined as its sister taxon, form a general pattern of the relationship within Coronaviridae.

### b. {*Alphacoronavirus*} clade (Figure 1, Figure S1 – S5, Table S1)

{*Alphacoronavirus*} clade includes all known species of the subgenus *Alphacoronavirus*.

Within this clade, all four trees showed that the following taxa are sisters:

a. *Minacovirus* 1 and 2 (Ferret coronavirus and Mink coronavirus 1);
b. *Pedacovirus* 1 and 2 (Porcine epidemic diarrhea virus and *Scotophilus* bat coronavirus 512);
c. *Setracovirus* 1 and 2 (Human coronavirus NL63 and NL63-related bat coronavirus BtKYNL63-9b);
d. *Decacovirus* 1 and 2 (Bat coronavirus HKU10 and *Rhinolophus ferrumequinum* alphacoronavirus HuB-2013);
e. *Minunacovirus* 1 and 2 (Miniopterus bat coronavirus 1 and Miniopterus bat coronavirus HKU8).

Thus, within {*Alphacoronavirus*} clade, we were able to find five smaller clades (subclades): {*Decacovirus*}, {*Minacovirus*}, {*Minunacovirus*}, {*Pedacovirus*} and {*Setracovirus*}with the exact correspondence of each of these clades to the previously established subgenera of the genus *Alphacoronavirus*.

From our results (Figure 1, Figure S1 – S5, Table S1) it is also clear, that

a. monotypic subgenus *Colacovirus* (Bat coronavirus CDPHE15) is a sister of *Pedacovirus* clade in all of the trees;
b. monotypic subgenus *Duvinacovirus* (Human coronavirus 229E) is a sister of the *Setracovirus* clade in all of the trees, and
c. monotypic subgenus *Tegacovirus* (Alphacoronavirus 1) is a sister of the *Minacovirus* clade in all of the trees.

The phylogenetic placement of monotypic subgenera *Luchacovirus* (Lucheng Rn rat coronavirus), *Myotacovirus (Myotis ricketti* alphacoronavirus Sax-2011) and *Rhinacovirus (Rhinolophus* bat coronavirus HKU2) depends on the method of the analyses (Figure 1, Figure S1 – S5, Table S1). It is worth stressing, however, that all of the cladistic methods (Figure S1 – S3), but not the ML method (Figure S4) have placed *Luchacovirus* as a sister of the {*Alphacoronavirus*}.

### c. {*Betacoronavirus*} clade (Figure 1, Figure S1 – S5, Table S1)

{*Betacoronavirus*} clade includes all known species of the subgenus *Betacoronavirus*.

All of the analyses argue in favor of the general simple hierarchical relationship ({*Embecovirus*} ({*Merbecovirus*} ({ *Nobecovirus*} (*Hibecovirus*, { *Sarbecovirus*})))) within *Betacoronavirus* clade (Figure 1, Figures S1 – S5).

The relationships of four species of the subgenus *Embecovirus* (1-4) (Betacoronavirus 1, China Rattus coronavirus HKU24, Human coronavirus HKU1 and Murine coronavirus), that formed a clade with the same name {*Embecovirus*}, the sister of {*Betacoronavirus*} clade, depend on the method of the analysis (Figure 1, Figures S1 – S5; Table 1).

All four species of subgenus *Merbecovirus* form a clade *{Merbecovirus*}. In all trees, the *Merbecovirus* 1 (Hedgehog coronavirus 1) is a sister to the clade that contains three other members of the subgenus *Merbecovirus*: namely, *Merbecovirus* 2 (Middle East respiratory syndrome-related coronavirus (MERS)), *Merbecovirus* 3 (*Pipistrellus* bat coronavirus HKU5), and the constant sister of the later *Merbecovirus* 4 (*Tylonycteris* bat coronavirus HKU4) (Figure 1, Figures S1 – S5; Table S1).

The {*Sarbecovirus*} clade that corresponds to the subgenus *Sarbecovirus* includes the viruses of severe acute respiratory syndrome-related coronavirus (SARS), the newly discovered monophyletic SARS-Cov-2 (two accessions have been included to the analyses, SARS-Cov-2a and SARS-Cov-2b), as well as CoV-ZC45 and SARS Cov ZS B.

Clade *{Sarbecovirus-SARS* plus SARS Cov ZS B} (SARS-Clade of the Figure 1) is a sister of the remaining species of *{Sarbecovirus*}.

Depending on the analysis, either bat coronaviruses RmYN02 or RaTG13 have been placed as a sister of SARS-Cov-2. The pangolin coronavirus (isolate MP789) has been defined as a sister of the clade {RmYN02 plus RaTG13 plus SARS-Cov-2} in all of the analyses.

All trees define the monotypic subgenus *Hibecovirus* (Bat Hp-betacoronavirus) as a sister of { *Sarbecovirus*} clade.

Two members of subgenus *Nobecovirus*, namely *Nobecovirus* 1 (*Rousettus* bat coronavirus GCCDC1) and *Nobecovirus* 1 (*Rousettus* bat coronavirus HKU9) are sister taxa (Figure 1, Figures S1 – S5; Table S1).

### d. {*Gammacoronavirus*} clade (Figure 1, Figure S1 – S5, Table S1)

Two subgenera of the genus *Gammacoronavirus*, namely subgenus *Cegacovirus* (with the single species Beluga whale coronavirus SW1) and subgenus *Igacovirus* (with a single species Avian coronavirus) formed a clade {*Gammacoronavirus*} in all of the analyses (Figure 1, Figures S1 – S5).

### e. {*Deltacoronavis*} clade (Figure 1, Figure S1 – S5, Table S1)

Five subgenera of genus *Deltacoronavis*, namely subgenus *Andecovirus* (with single species Wigeon coronavirus HKU20), subgenus *Buldecovirus* (1-4) (with four species: Bulbul coronavirus HKU11, Coronavirus HKU15, Munia coronavirus HKU13 and White-eye coronavirus HKU16), subgenus *Herdecovirus* (with one species Night heron coronavirus HKU19) and subgenus *Moordecovirus* (with single species Common moorhen coronavirus HKU21), have formed the *{Deltacoronavis}* clade in all of the analyzes. Also, all of the analyses argue in favor of the simplest hierarchy of the relationships within this clade: (*Andecovirus (Herdecovirus* (*Moordecovirus* ({*Buldecovirus*})))). Within {*Buldecovirus*} clade *Buldecovirus* 2 (Coronavirus HKU15) and *Buldecovirus* 3 (Munia coronavirus HKU13) appeared to be sisters in all of the analyses, the relationships between the other members of the { *Buldecovirus*} clade depends on the method of the analysis. Monotypic subgenus *Andecovirus* is a sister of *{Deltacoronavis}* clade.

### f. On monophyly of non-monotypic taxa of Coronaviridae (Figure 1, Figure S1 – S5, Table S1)

As is perhaps clear from above, all four trees show that all four genera of subfamily Orthocoronavirinae (family Coronaviridae), namely *Alphacoronavirus*, *Betacoronavirus*, *Deltacoronavirus*, and *Gammacoronavirus*, are monophyletic (Figure 1, Figures S1 – S5).

All non-monotypic subgenera of all four genera of subfamily Orthocoronavirinae (family Coronaviridae), namely subgenus *Decacovirus* (genus *Alphacoronavirus*), subgenus *Minacovirus* (genus *Alphacoronavirus*), subgenus *Minunacovirus* (genus *Alphacoronavirus*), subgenus *Pedacovirus* (genus *Alphacoronavirus*), subgenus *Setracovirus* (genus *Alphacoronavirus*), subgenus *Embecovirus* (genus *Betacoronavirus*), subgenus *Merbecovirus* (genus *Betacoronavirus*), subgenus *Nobecovirus* (genus *Betacoronavirus*), and subgenus *Buldecovirus* (genus *Deltacoronavirus*) are monophyletic in all of the analyses (Figure 1, Figures S1 – S5).

## Discussion

Several phylogenies of Coronaviridae (or parts thereof) have been published; however almost all of these remain unrooted (for example (*2, 3, 6, 8–11*) among others), or if the phylogenetic tree occasionally appears as rooted (for instance (*4, 7*)) the taxonomic sampling of such studies remain incomplete or even arbitrary. We would like to stress that this study seeks to produce comprehensive rooted phylogenetic Coronaviridae trees and that the analyses herein have used the best taxonomic summaries of family Coronaviridae currently established by ICTV (2019) (*1–4*).

### a. Rigorous rooted trees validate tested proposals of the relationships within Coronaviridae

Rooted trees produced by the four methods herein allow for the effective testing of various proposals regarding the relationships of viruses within the Coronaviridae family.

1. Our results argue in favor of Bukhari *et al*. (*2*) proposal regarding the new family Abyssoviridae of the order Nidovirales (current monotypic subfamily Letovirinae with subgenus *Milecovirus*) (Table S1), which was unfortunately based on an unrooted phylogenetic tree and limited taxonomic sampling. After Bukhari *et al*. (*2*) we also found that two subfamilies of Coronaviridae, namely Letovirinae (family Abyssoviridae in Bukhari *et al*. (*2*)) and Orthocoronavirinae are sisters (Figure 1, Figures S1 – S5). This solution is consistent with the familial rank of both taxa (*2*). We would recommend to accept the monotypic subfamily Letovirinae at the familiar rank (*2*).
2. Tokarz *et al*. (4) proposed the (((A, B) Γ) Δ) hypothesis of the general pattern of relationship within subfamily Orthocoronavirinae; however this was again based on limited taxonomic sampling and an unrooted network. Our results clearly argue in favor of this hypothesis. Keeping in mind the constant placement of subgenus *Milecovirus* (MLeV), we can extend this pattern to the relationship ((((A, B) Γ) Δ) MLeV). This natural hierarchy within Coronaviridae is in principle congruent to the general pattern of their hosts: (((Mammals) Birds plus Mammals) Amphibia) (Table S1).
3. Subgenus *Hibecovirus* has been proposed as a sister taxon of the subgenus *Sarbecovirus* (reviewed in (*5*)). This proposal, which again was made on the basis of unrooted trees, also benefited from reexamination using the comprehensive rooted trees produced in this study. All four methods validate the sister relationship of the subgenera *Hibecovirus* and *Sarbecovirus* clade (Figure 1, Figures S1 – S5).
4. Because of our particular interest in regards to SARS-Cov-2, we would like to stress that all four methods have placed pangolin coronavirus as a sister of the clade {RaTG13 plus RmYN02 plus SARS-Cov-2} (Figure 1, Figures S1 – S5), confirming that bat coronaviruses RaTG13 or RmYN02, but not the pangolin coronavirus, are indeed the closest relatives of SARS-Cov-2 (Figure 1, Figures S1 – S5). In other words, contrary to some recent suggestions (reviewed in *17, 18*) the rooted trees produced herein confirm that either bat coronavirus RaTG13 (*21*) or RmYN02 (*24*), but not the pangolin coronavirus, is an immediate sister of SARS-Cov-2 (Figure 1, Figures S1 – S5).

Differences in details within clade relationships do exist between the ML method and each of the additional cladistic methods newly applied to molecular sequence data of Coronaviridae. For example the MP, AC and 3TA trees (Figures S1–S3), but not ML trees (Figure S4), have placed Lucheng Rn rat coronavirus (genus *Alphacoronavirus*, subgenus *Luchacovirus*) as a sister taxon to the clade that contains all of the remaining members of genus *Alphacoronavirus*.

Similarly, both conventional phylogenetic methods (MP and ML) have defined bat coronavirus RmYN02 as a weakly supported sister of the SARS-Cov-2 (Figures S1, S4). However both Hennigian methods (3TA and AC) are, in contrast, placed bat coronavirus RaTG13, but not RmYN02, as a sister of this newly discovered coronavirus (Figures S2, S3).

All four analyses are initially based on a common molecular matrix (22,489 bp G-Block version of the 47-genomes alignment) (Supplementary Materials). Differences between analyses are likely a result of how each method deals with conflict to form the optimal trees. Nevertheless, the similarity between the tree topologies suggests that, regardless of method, many of the nodes are ‘true’ summaries of the data and that the data themselves are relatively noise-free (Figure 1, Figures S1 –S5).

### b. On relationships and hosting of newly discovered *Sarbecovirus* SARS-Cov-2

Even from the elementary comparative point of view, it is clear that every Coronaviridae virus seems to be well defined and well separated from the others sometimes by hundreds or (more commonly) thousands of single nucleotide positions (SNPs) (Table S2, S3). For example, the minimal relationship {RmYN02 {RaTG13 plus SARS-Cov-2}} (Figures S2, S3), based on the 29,907 bp complete genomic alignment of three taxa (RmYN02, RaTG13 and SARS-Cov-2 (MN908947)), implies 1,329 informative SNPs (or, respectively, 1,329 3TSs) from the total 2,467 variable characters.

Thus, even closely related viruses from the *Sarbecovirus* clade (including the newly discovered SARS-Cov-2 and bat coronaviruses RaTG13 and RmYN02), are all remarkably different from one another from a comparative standpoint. The same is true for every relationship within Coronaviridae we have recovered in our analyses.

Such simple observations automatically exclude the possibility of the recombination origins of SARS-Cov-2 (reviewed in (*19*), see also (*8, 20*)) as well as other similar propositions. Based on the data available to date, including the comprehensive trees produced herein (Figure 1, Figures S1 – S5), the focus should shift to the static aspect of the problem.

The monophyly of all the genera of Coronaviridae, as well as all of its non-monotypic subgenera, and also the general relationship ((((A, B) Γ) Δ) within the subfamily Orthocoronavirinaeis, can also be demonstrated within the pure comparative analytical framework (Figures S2 and S3). Within the later, no member of *Sarbecovirus* clade is an ancestor of SARS-Cov-2. This static view may be critical in discussing the general simple pattern in hosting of SARS-Cov-2.

Recent studies of SARS-related coronaviruses have suggested that bats harbor close relatives to SARS-Cov-2 (for example (*21*)), and that pangolins may be natural hosts of this member of genus *Betacoronavirus* (*8, 18, 26*), leading to the hypothesis of animal to human transmission of SARS-Cov-2 (“The presence in pangolins of an RBD very similar to that of SARS-CoV-2 means that we can infer this was also probably in the virus that jumped to humans” (*18*)).

However, the search of other hosts, as well as the related exotic ways of transmission of SARS-Cov-2 from these hypothetical hosts to humans, is based on a set of the complicated assumptions (for example, “The sequence similarity in the spike receptor binding domain between SARS-CoV-2 and a sequence from pangolin is probably due to an ancient intergenomic introgression…” (*26*)) and also ignores the simple possibility of the original human-based hosting of SARS-Cov-2.

In fact, the hosting of the viruses that are genetically related to SARS-Cov-2 by bats or pangolins is not, strictly speaking, an argument in favor of the animal hosting of SARS-Cov-2, especially because the latter virus had never been detected inside of animals such as bats. The clear possibility of the original human hosting of SARS-Cov-2, unfortunately, had not been discussed in scientific literature. However, the extremely high contagiousness of SARS-Cov-2 is almost improbable if the virus had been transmitted from animals to humans just several months ago. Alternatively, it is possible that a form of SARS-Cov-2 may have been existed in some parts of the human population before the current pandemic, and that, perhaps, the virus had been effectively suppressed by the human immune system for a sometime prior. Seven human coronaviruses have been identified to date (reviewed in (*22*)), yet at least three of these namely, Human coronaviruses OC43 and HKU1 from subgenus *Embecovirus* and NL63 from subgenus *Setracovirus* (Table S1) are globally distributed viruses where no animal hosts have been proposed.

With the information available to date, there is no direct evidence to suggest which hypothesis, animal to human transmission (either the recent or less recent one (*25, 26*)) or original human hosting, is true. The evidence only tells us how viruses are related within a hierarchical classification (Figure 1).

### c. On taxonomy, naming, and nomenclature of the coronaviruses

Because the taxonomy and nomenclature of the viruses are still “under construction” (*13–16*), the names of the virus species frequently remain non-binary, even within the taxonomic statements of ICTV (*13–16*). Simultaneously, the current circumscriptions of the two largest genera of the Coronaviridae (*Alphacoronaviruses* and *Betacoronaviruses*) are very complicated. For example, current genus *Alphacoronavirus* currently circumscribes 12 subgenera and current genus *Betacoronavirus* circumscribes five subgenera. These two observations cause issues with a clear naming of viruses on the phylogenetic trees of Coronaviridae, as well as with the reading of these trees, especially by a non-specialist.

Here we resolved these issues by using the ICTV summaries (*13–16*) of the **subgeneric** names of Coronaviridae as the basic units of our notation where possible in the analyses and trees. In other words, whenever possible, abbreviations and trivial names were avoided to improve clarity. The recently discovered SARS-Cov-2 virus and a few related viruses are the exceptions (Table S1).

Reconsideration of the current circumscriptions of Coronaviridae genera may provide a simpler and more informative taxonomic system of both naming and nomenclature. Accepting traditional genera of the family at the rank of a tribe and, simultaneously, the current subgenera (all of which are monophyletic (Figure 1, Figures S1 – S5)) at the generic rank seems to be the easiest heuristic way to incorporate the Linnaean principles of the binary nomenclature right to the classification of the family. For example, the current member of the subgenus *Sarbecovirus* (current genus *Betacoronavirus*) virus SARS-Cov-2 may be easily named ***Sarbecovirus* species, abbreviation: sp. (e.g.: *Sarbecovirus* sp.)**, where “species” (sp.) is any available epithet. Numerous currently discovered variants of it (summarized in (*6*)) can be *in principle* established as forms of varieties of the same species. The trivial name “severe acute respiratory syndrome-related coronavirus 2” and correspondent abbreviation “SARS-Cov-2” can be simply listed in the description of the species.

It is also critical to consistently involve the type method of biological taxonomy to the taxonomy of the viruses. As a simple example, we would like to suggest that the ICTV approved GenBank Accession number (ideally the reference to the whole genome of the virus) can be used as a nomenclature type of any virus species. For example the GenBank Accession number MN908947 can be treated as a nomenclature type of newly described *Sarbecovirus* (SARS-Cov-2). Such a number implies the name/abbreviation of the biological isolate as well as other useful information. The higher nomenclature categories (tribes, genera, families etc.) can be typified by the names of the species (genera etc.), exactly in a manner of the plant or animal names. The nomenclature classification of plants and animals has developed over hundreds of years, and as such is robust and well tested. Adopting the Linnaean binary nomenclature for viruses will increase the universality of the system, and thereby lead to more consistent information content and information exchange.

We believe that such simple recommendations are fully consistent with the principles of the future binary nomenclature of viruses, currently summarized by Siddell *et al. (15, 16*) and others (*23*) and may be useful for the different families of viruses. Such a nomenclature system would help scientists find information about a particular taxon easily and quickly, which is a high priority when accurate and timely identification is required during pandemic outbreaks.

## Author contributions

E.V.M.: Conceptualization, Methodology, Software, Validation, Formal Analysis, Investigation, Resources, Data Curation, Writing - original draft, Writing - review & editing, Visualization, Supervision, Project administration. N.E.M.: Conceptualization, Methodology, Software, Formal Analysis, Investigation, Resources, Writing - review & editing, Visualization. M.C.E.: Conceptualization, Methodology, Formal Analysis, Investigation, Writing - original draft, Writing - review & editing, Supervision. D.M.W.: Conceptualization, Methodology, Formal Analysis, Investigation, Writing - original draft, Writing - review & editing, Supervision. M.L.T.: Conceptualization, Investigation, Data Curation, Writing - original draft, Writing - review & editing, Supervision.

## Data and materials availability

“All data is available in the main text or the supplementary materials.”

## Supplementary Materials

### This PDF file includes

Materials and Methods

Supplementary Text

Figures S1 to S5

Captions for Figures S1 to S5

### Other Supplementary Materials for this manuscript include the following

Tables S1 to S3

Captions for Tables S1 to S3

## Materials and Methods

### a. Taxonomic sampling of the study

Thirty nine ICTV-approved genomes (*1–4*) of all species of Coronaviridae have been used in this study (Table S1).

Additionally to these 39 genomes, we also included along with the final alignments

a. the published genome MN908947 of SARS-Cov-2(*5, 6*) (Table S1);
b. an unpublished genome of the same virus species (MN988713, Tao *et al*., unpublished) (Table S1);
c. the genome of bat SARS-like coronavirus (bat-SL-CoV-ZC45; MG772933) (Table S1) that has been previously used in other comparisons to SARS-Cov-2 (for instance (*5–7* and others));
d. - f. bat coronavirus RaTG13 (MN996532 (*8*) (Table S1)), pangolin coronavirus (MT121216 (*9*)) (Table S1) and RmYN02 (GISAID: EPI_ISL_412977) (*32*) (Table S1)) as they have previously been proposed as the closest known relatives of SARS-Cov-2 (reviewed in (*9, 10, 32, 33*));
g. an additional SARS related genome ZS-B (AY394996) (Table S1).

The genome of recently discovered *Microhyla letovirus* 1 or MLeV virus (subgenus *Milecovirus*) (*11*) was also included with the alignments and analyses. This genome was available upon courtesy request from Prof. B. W. Neuman (Texas A&M University-Texarkana, TX, US) (Table S1).

Based on the summary of Maclachlan *et al*. (*12*), genus *Torovirus* (family Tobaniviridae, order Nidovirales) was assumed as the best outgroup taxon of Coronaviridae, and ICTV-approved genome of *Torovirus* (AY427798) (Table S1) has been selected for this study.

The names of the major obtained relationships on all trees (Figure S1–S5) excluding the names of the families and subfamilies, are written in italics due to the strong congruence with different taxonomic entities. Depending on the context, the names of the clades are provided in regular and/or curved brackets.

As the utility of phylogenetic trees depends on their clarity, the use of abbreviations and trivial names of viruses has been avoided whenever possible.

### b. Matrices and trees

All genomic alignments have been performed using MAFFT (*13, 14, 15*) following FFT-NS-I strategy with the command: mafft --inputorder --adjustdirection -- anysymbol --kimura 1 --maxiterate 1000 --6merpair input.

Poorly aligned positions have been removed from

1. the genomic alignment of Coronaviridae plus *Torovirus* and
2. the genomic alignment of subfamily Orthocoronavirinae with no outgroups included

using the program G-block (*16*) as implemented in SeaView (*17*) under the conditions of “less stringent” strategy of the algorithm (*16, 17*).

The G-block version of the genomic alignment of Coronaviridae plus *Torovirus* was also established as a binary matrix using simple “presence – absence” coding (reviewed in (*18*)) with the future inclusion of the “all-plesiomorphic” (“all-zero”) artificial taxon. In short, G-block based genomic alignment of the family Coronaviridae plus *Torovirus* was rewritten as a binary (01) matrix, where “zero” means “the absence of a nucleotide in this particular position of the alignment”, and “one” means “the presence of the nucleotide in this particular position of the same alignment”. For example, if the character-state of the character number 253 is equal to A (Adenine), than this can be written as “1000”, where “1” means “the A is present in position 253”, and “0” indicates that U(T), G and C are simultaneously absent on the same position.

Assuming that the “absence of the nucleotide” (the character-state “zero”) is a plesiomorphic character-state, we can add to the binary matrix “all-plesiomorphic” or “all-zeros” outgroup. The binary matrix with an “all-zeros” outgroup added was later used as an input into the script Forester v. 1.0 (*19*) following the command ruby trees.rb path_to_matrix_file (*19*) with future selection of the “Additional” forest of the maximal trees (relationships) (*19*) for Average Consensus (AC) analysis (*19–21*).

Manipulations with either the molecular or binary matrices and the tree-files have been performed with Mesquite v. 3.51 (*22*), PAUP* v. 4.0a (*23*) and FigTree v. 1.4.2 (*24*).

### 3. Analyses

The G-block version of the molecular alignment of Coronaviridae plus *Torovirus* was analysed by the standard Maximum Parsimony (MP) approach (Fitch Parsimony, reviewed in (*18*)), and by the three-taxon statement analysis (3TA) with fractional weighting (reviewed in (*18, 27, 28*) and implemented in Mavrodiev and Madorsky (*29*)).

Following the logic of Williams-Siebert (WS) representation of the unordered multistate data (reviewed in (*29*)) the three-taxon statement (3TS) permutations of the G-block-based alignment of Coronaviridae plus *Torovirus* were conducted with TAXODIUM version 1.2 using the command: taxodium.exe input_file_name.csv –idna –ob –og –fw –nex (*29*) taking values of the operational outgroup as equal to the values of *Torovirus*. All MP analyses have been performed in PAUP* (*23*) as in Mavrodiev *et al*. and Mavrodiev and Madorsky (*19, 29*). The resulted most parsimonious tree resulted standard MP analysis was *a posteriori* rooted relative to *Torovirus*.

The AC (*20, 21*) of the array of maximal trees was calculated using the program Clann version 4.1.5 (*25, 26*) as described in Mavrodiev *et al. (19*). The distance optimality criterion for the AC analysis was specified as a “distance with non-weighted least squares” (*20, 21, 23, 25*).

Following conclusions of Zhou, X. *et al*. (*30*), the Maximum Likelihood (ML) analysis of G-block alignment of Coronaviridae plus *Torovirus* was conducted with W-IQ-TREE (*31*) with implemented automatic model selection procedure (*31*). The resulted most probable tree was *a posteriori* rooted relative to *Torovirus*.

The MP Bootstrap support (BS) values have been calculated as described in (*29*) and (*31*). In the case of the ML analysis, the aLRT support values have been calculated instead of the ML BS supports, as implemented W-IQ-TREE (*31*).

The simplest “total” character differences between the G-block modified aligned genome sequences of subfamily Orthocoronavirinae (Table S2), as well as between three aligned genomes of bat coronaviruses RmYN02, RaTG13 and severe acute respiratory syndrome-related coronavirus 2 (SARS-Cov-2) (MN908947) with no aligned positions excluded (Table S3), was calculated in PAUP*(*23*) under the default options.

### d. Supplementary Text

The genomic alignment of Coronaviridae + *Torovirus* (outgroup) consists of 52,990 molecular characters (base pairs or bp), the G-block version of this alignment is of 22,489 characters with 19,550 of those are parsimony-informative. This alignment was the target of future MP, 3TA, AC and ML analyses.

The genomic alignment of subfamily Orthocoronavirinae with no outgrop included is of 49,881 bp. The G-block version of this alignment consists of 23,431 bp. Later, the alignment has been used only to calculate the simplest pairwise distances (“total character differences”) between all of the members of the subfamily included in the analyses (Table S2). The genomic alignment of bat coronaviruses RmYN02, RaTG13 and severe acute respiratory syndrome-related coronavirus 2 (SARS-Cov-2) (MN908947) is of 29,907 molecular characters. This alignment with no characters excluded was used to calculate the total character differences (Table S3) between newly discovered SARS-Cov-2 and two of its closest relatives. Such a measure is the simplest expression of the pairwise distance between aligned molecular sequences and indicates solely the total number of different single nucleotide positions (or SNPs) between them. For example, the number 1138 in the Table S3 means that the aligned genomes of human coronavirus SARS Cov 2 (accession “a”) and the bat coronavirus RaTG13 are different from each other by 1138 SNPs. For instance, in position # 37 of the same molecular alignment the value of SARS Cov 2 is equal to “C” and the value of RaTG13 is equal to “G”. The total number of such SNPs equals 1138.

The standard MP analysis of the 22,489 bp G-block alignment resulted in the single most parsimonious tree of 208,176 steps (CI = 0.2505, RI = 0.4954) (Figure S1).

The 3TS representation of the same 22,489 bp G-block alignment resulted 39,621,820 3TSs (binary characters), all parsimony-informative and fractionally weighted, with the most parsimonious fit of 20,786,459.7424 steps (RI = 0.5706) (Figure S2).

The presence-absence re-coding of the genomic alignment of Coronaviridae + *Torovirus* resulted in a matrix of 73,258 binary characters from which 65,435 characters can be established as a rooted trees (maximal relationships). The AC analyses of the forest of these 65,435 relationships resulted in a single AC tree of the score 0.00911 (Figure S3).

For ML analysis of the 22,489 bp G-block matrix of Coronaviridae and outgroup (*Torovirus*), GTR+F+R10 model has been automatically selected by W-IQ-TREE as a best-fit model based on either corrected and non-corrected Akaike Information Criteria, as well as on Bayesian Information Criterion. The resulted single most probable (ML) tree has the best score (log likelihood) equal to −766940.2344 (Figure S4).

## Captions for Figures S1 to S5

**Figure S1.**
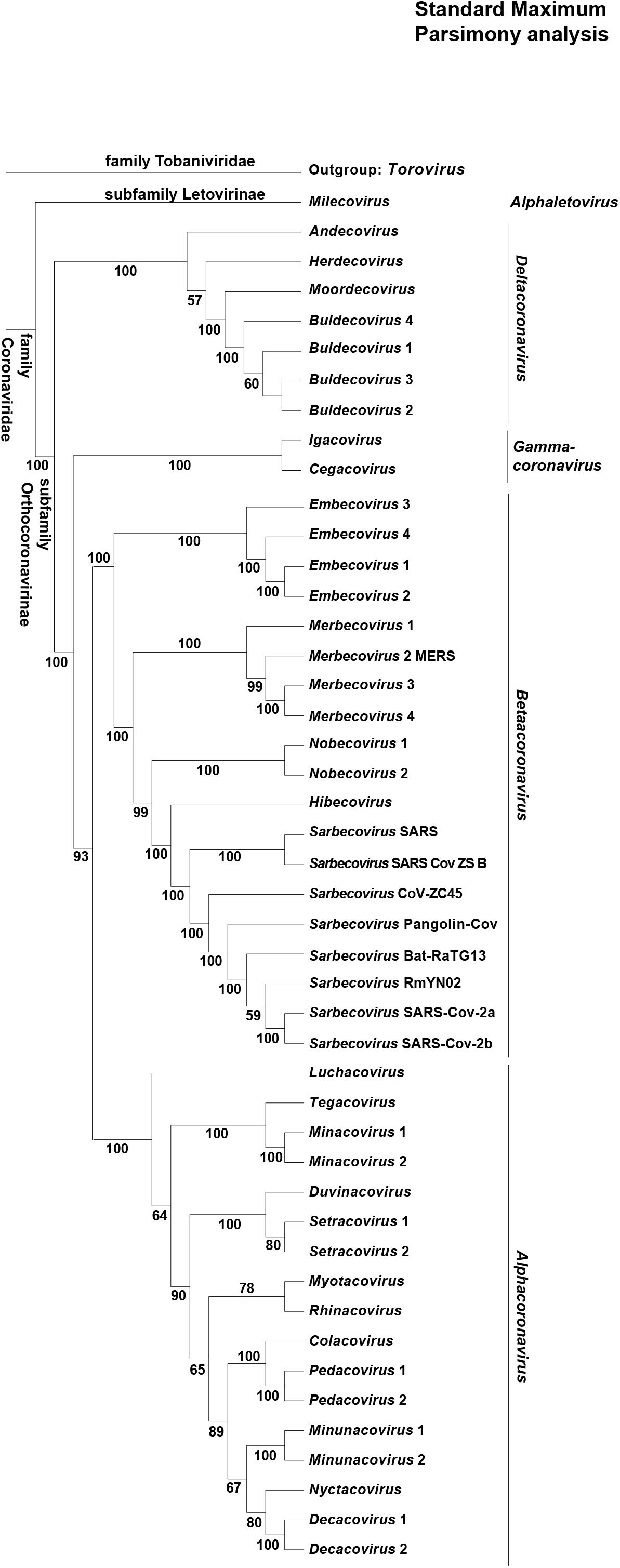
Single most parsimonious tree recovered from the standard MP analysis (Fitch Parsimony) of the 22,489 bp genomic alignment of Coronaviridae + *Torovirus*. Tree was *a posteriori* rooted relative to *Torovirus*.

**Figure S2.**
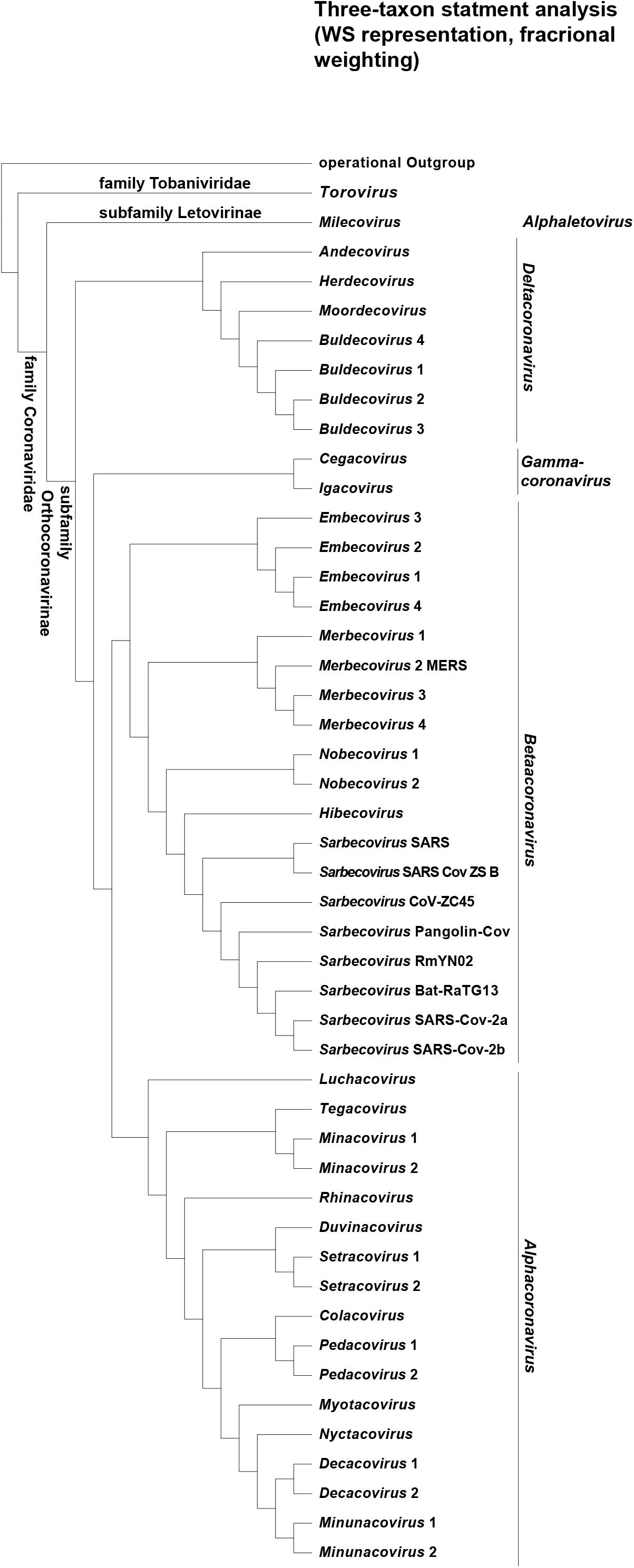
Most parsimonious hierarchy of patterns recovered from MP analysis (Wagner parsimony) of 39,621,820 3TS WS representation of the 22,489 bp genomic alignment of Coronaviridae + *Torovirus*. The values of the operational outgroup were fixed as values of *Torovirus*.

**Figure S3.**
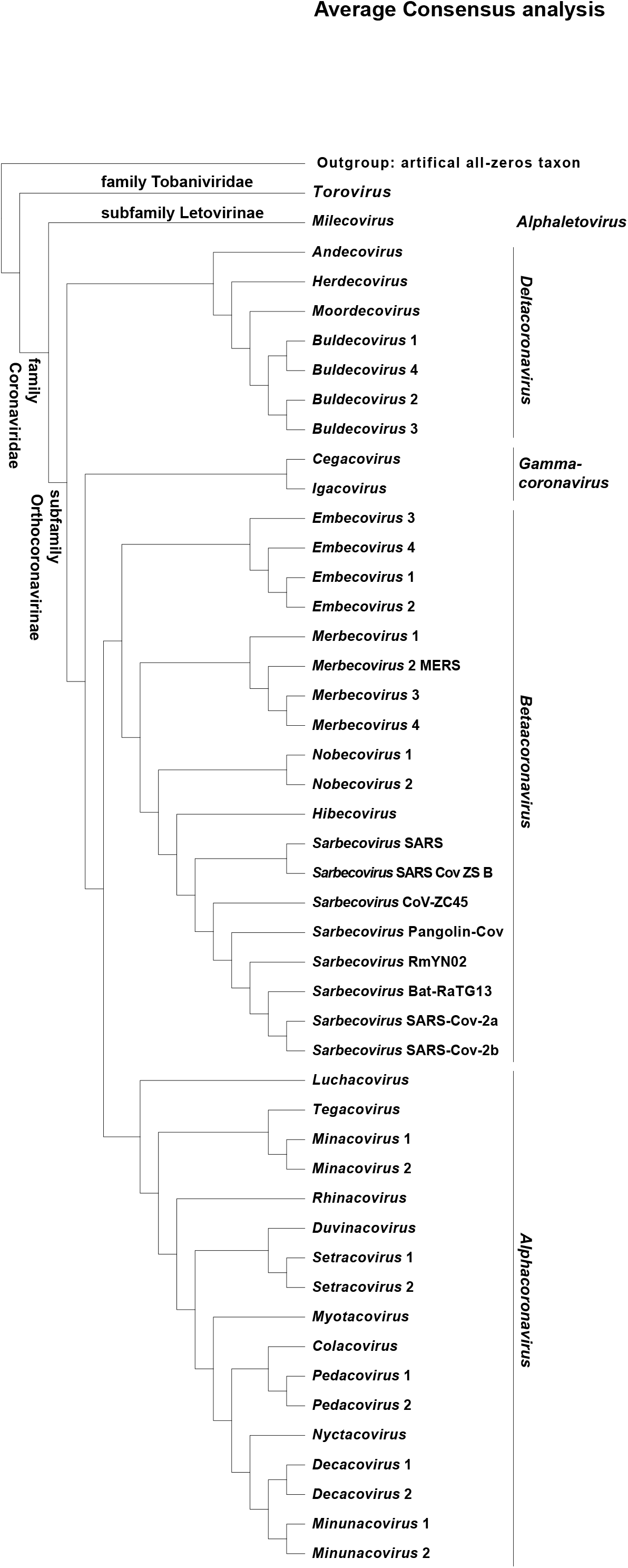
The average consensus tree from the analysis of the forest of 65,435 rooted trees (maximal relationships) derived from the “presence-absence” representation of the original 22,489 bp genomic alignment of Coronaviridae + *Torovirus*.

**Figure S4.**
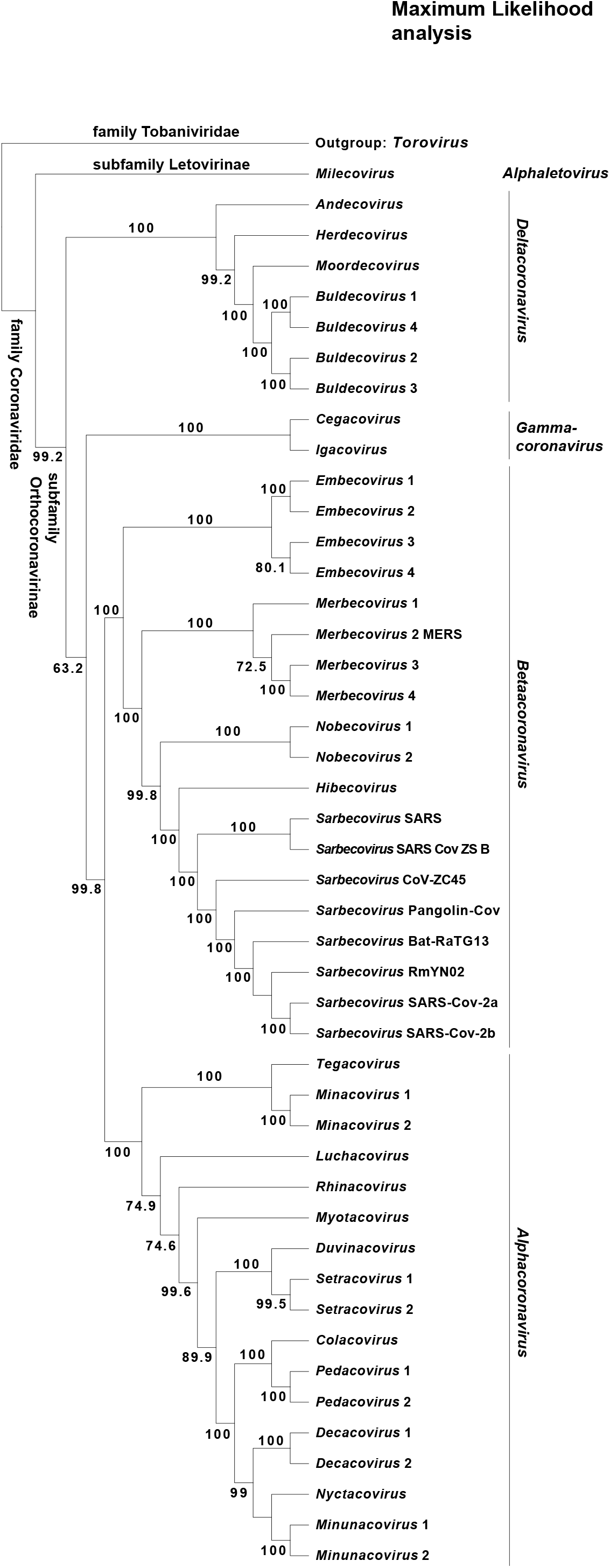
Most probable tree recovered from the ML analysis of the 22,489 bp genomic alignment of Coronaviridae + *Torovirus*. Tree was *a posteriori* rooted relative to *Torovirus*.

**Figure S5.**
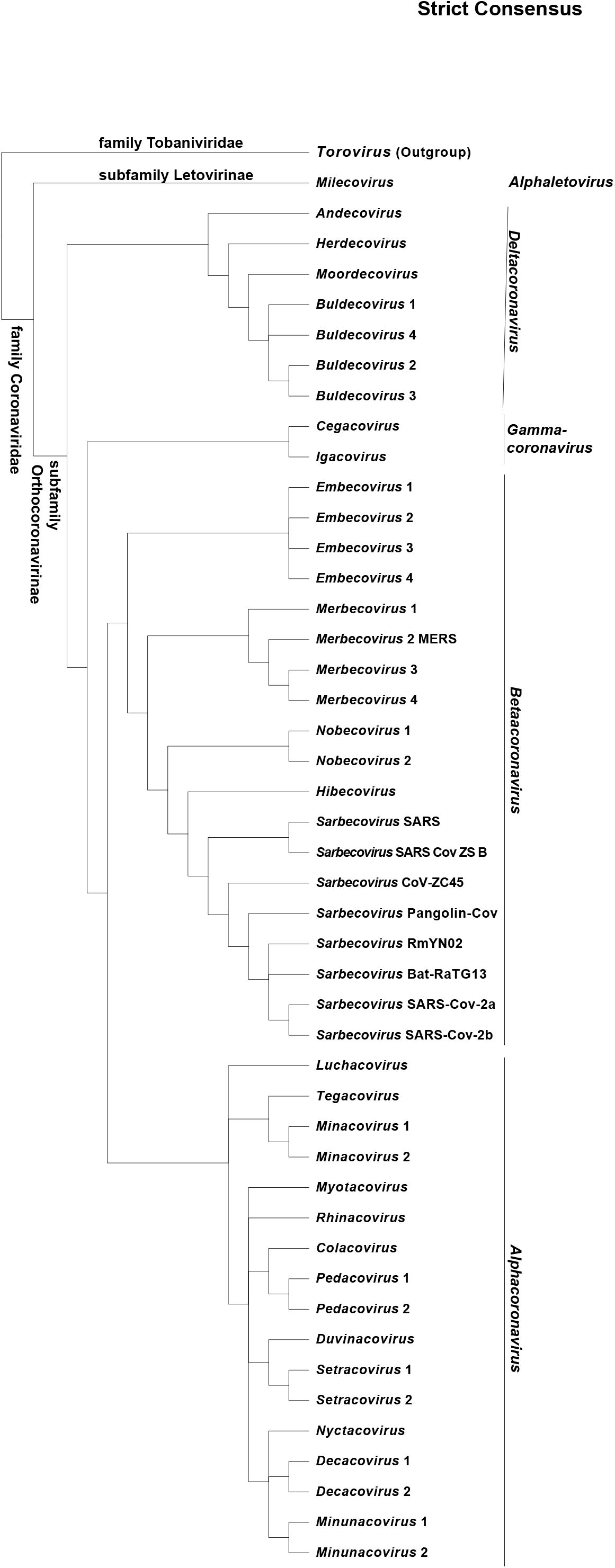
Strict Consensus of four trees produced by three different methods of cladistic analysis as well as by Maximum Likelihood method using modified genomic alignment of Coronaviridae + *Torovirus* (Figures S1 – S4).

## Captions for Tables S1 to S3

**Table S1.**
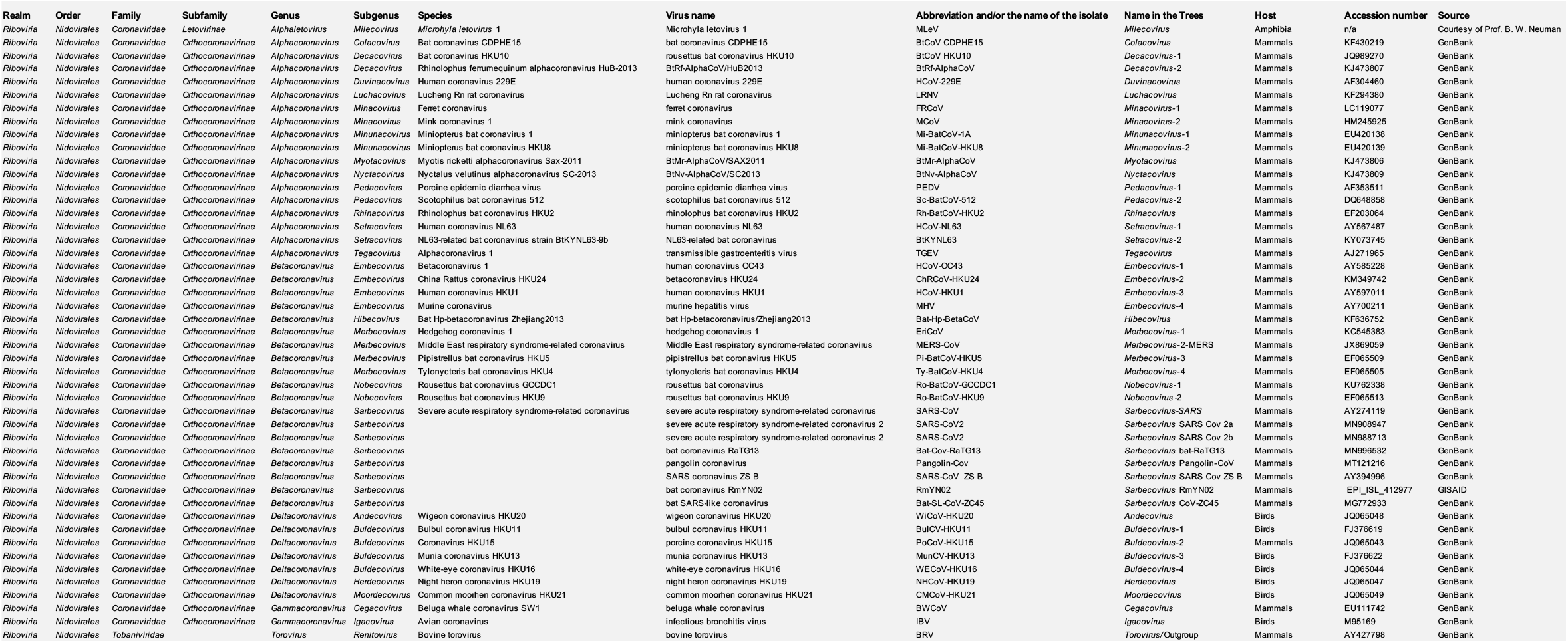
Taxonomic sampling of the study and related data.

**Table S2.**
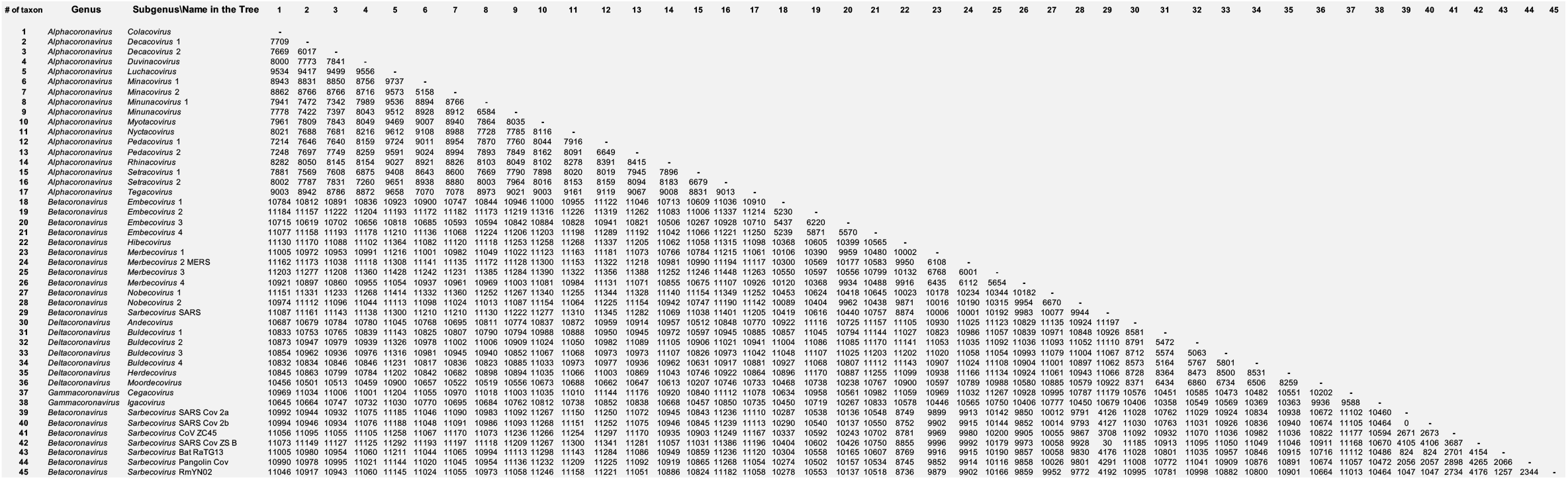
Total character differences between the aligned genomes of the all coronaviruses from the subfamily Orthocoronavirinae (family Coronaviridae, order Nidovirales) based on the G-block version of the genomic alignment.

**Table S3.** Total character differences between the aligned genomes of severe acute respiratory syndrome-related coronavirus 2 (SARS-Cov-2) (MN908947) and bat coronaviruses RmYN02 and RaTG13 (family Coronaviridae, subfamily Orthocoronavirinae, genus *Betacoronavirus*, subgenus *Sarbecovirus*).

| # of taxon | Genus | Subgenus\Name in the Tree | 1 | 2 | 3 |
| --- | --- | --- | --- | --- | --- |
| 1 | *Betacoronavirus* | *Sarbecovirus* SARS Cov 2a | - |  |  |
| 2 | *Betacoronavirus* | *Sarbecovirus* Bat RaTG13 | 1138 | - |  |
| 3 | *Betacoronavirus* | *Sarbecovirus* RmYN02 | 1803 | 2032 | - |

